# Effect of simulated microgravity on gene expression during embryogenesis of *Arabidopsis thaliana*

**DOI:** 10.1101/471037

**Authors:** Michaela Švécarová, Markéta Kovalová, Vladan Ondřej

## Abstract

Gravitational force is one of environmental factors that influence growth and development of plants. Changes in this force, including microgravity, can be one of the stress factor which plants have to adapt to cope with. That kind of stress can lead to several abnormalities such as chromosomal breakage, morphological abnormalities or changes in gene expression. The aim of this study was to examine the influence of simulated microgravity on gene expression of *Arabidopsis thaliana* embryos by use Random Positioning Machine (RPM). RPM is laboratory facility that can generate conditions comparable to the true microgravity. This paper studies effect of simulated microgravity on expression of genes which are involved in plant embryogenesis (*LEAFY COTYLEDON, LATE EMBRYOGENESIS ABUNDANT*), antioxidative system (*CATALASE*), mechanical stimuli (*TOUCH*) and gravitropism (*SCARECROW, SHOOT GRAVITROPISM2*). Changes in gene expression were detected using quantitative real-time PCR (qRT-PCR). Several of tested genes had increased transcript levels after the influence of simulated microgravity. Specifically, catalase (*CAT3*), LECs (*LEC1*), touch (*TCH2/CML24*), and gravitropism (*SGR2*) genes achieved significantly increased relative expression (level ≥ 2). The changes in the levels of expression on embryos of *Arabidopsis* depend on the type of genes and principally on the timing of the influence of the simulated microgravity.

## Introduction

Microgravity has been demonstrated to have unavoidable impacts on living organisms during space flights and can be considered as important factor for estimating potential health risks for astronauts. Changes in the strength of the gravitational field are most surely yet another type of stress, which is transduced by signaling chains (Martzivanou *et al*., 2006). In order to study the effects of simulated microgravity, we utilized a random positioning machine (RPM) in the laboratory. A RPM is a version of 3D clinostat, which continuously realizes random changes in the orientation relative to the gravity vector in a biological experiment (Borst *et al*., 2009). The first experiment of *Arabidopsis thaliana* in microgravity was performed by Merkys and Laurinavicius (1983), they obtained some viable seeds, on the other hand several of seed contained nonviable embryos (reviewed in Link *et al*., 2014). Simulated (micro)gravity is stressful for plants, and has an influence on plant growth, as well as on cellular and molecular responses such as cell cycle, embryogenesis, seeds, photosynthesis, gravitropic sensing/response, cell wall composition/properties, and changes of gene expression (Link *et al*., 2014). The growth and development of plants is highly influenced by various external factors including abiotic and biotic stresses. The antioxidant system is involved during plant stress and is comprised of several enzymes. One of them is catalase (*CAT*, H_2_O_2_:H_2_O_2_ oxidoreductase; EC 1.11.1.6), which is not plant specific but is present in almost all living organism (Islam *et al.*, 2015). In *Arabidopsis* genome are located three catalase genes (*CAT1, CAT2* and *CAT3*). These genes encode individual subunits, which associate to form at least six isozymes. *CAT1* and *CAT3* genes are localized on chromosome 2, and *CAT2* is localized on chromosome 4 (McClung, 1997; Mhamdi *et al*., 2010). The study (Alam and Ghosh, 2018) described that the expression of *CAT1* was found in floral organs and seeds. In photosynthetic tissues, also in the seeds and roots is primarily expressed *CAT2* gene and it is known, that gene *CAT3* is associated with vascular tissues, but also with the leaves (Mhamdi *et al*., 2010). The researchers (Frugoli *et al*., 1996) say that in mature *Arabidopsis* rosettes are detected all three transcripts of *CATs*; however, *CAT2* and *CAT3* transcripts are much more plentiful than those of *CAT1*. The expression of *CAT2* and *CAT3* in *Arabidopsis* is controlled by circadian rhythms, with a morning-specific phase for *CAT2* and an evening-specific phase for *CAT3* (Du *et al*., 2008; McClung *et al*., 1997). In addition to that, *CAT2* revealed downregulation during leaf senescence, but senescence and also age induced the expression of *CAT3* (Zimmermann *et al*., 2006).

Other *Arabidopsis* genes examined in this paper are the *LEAFY COTYLEDON* (*LEC*) genes, specifically: *LEC1*, *LEC2*, and *FUS3* (*FUSCA3*), which possess principal main specificity on embryo development (Gaj *et al*., 2005; Harada *et al*., 2001). *LEC* genes are unique in that they are required for normal development during both the morphogenesis and maturation phases. For specificity suspensor cell fate and cotyledon identity are required *LEC* genes in early embryogenesis (Harada *et al*., 2010). The authors (Stone *et al*., 2001) displayed that these mentioned genes are also necessary in late embryogenesis during the maturation phase for the acquisition of desiccation tolerance. From this finding, we can say that *LEC* TFs are appropriate and essential for induction of the maturation phase of zygotic embryogenesis. Direct targets of *LECs* TF are genes involved in storage macromolecule synthesis (Braybrook and Harada, 2008). The author (Harada, 2001) described, how *LEC* genes play important roles during maturation. Also ectopic expression of *LEC1* and *LEC2* can induce embryo development in vegetative cells (Gaj *et al*., 2005; Stone *et al*., 2001).

Another gene studied, the LATE EMBRYOGENESIS ABUNDANT (*LEA*) gene, is involved during embryogenesis. LEA proteins are present in plants, and they accumulate to high levels during the developmentally regulated period of dehydration at the end of seed development and also in the vegetative organs, suggesting a protective role during times of limited water (Battaglia *et al*., 2008). The mentioned proteins were not plant specific, but were discovered in phylogenetically distant organisms and have always been connected to abiotic stress tolerance, particularly desiccation tolerance (Hundertmark and Hincha, 2008). According to their sequence similarities, the LEA proteins are divided into seven groups (Battaglia, 2008). In *Arabidopsis*, 51 LEA protein encoding genes in the genome have been identified; 22 of them are highly expressed in non-seed tissues. The value of the expression of *LEA* is enhanced primarily by cold, drought, and salt treatment. On the contrary, only one *LEA* gene’s expression is enhanced by heat (Hundertmark and Hincha, 2008).

The other studied genes of this paper in *Arabidopsis thaliana* were *TCH* genes, which were the first to be described as being upregulated in responses to mechanical stress, being induced within 30 min after mechanical stimuli. This investigation is reviewed by Braam (2005). In Arabidopsis, *TCH1* encodes calmodulin (*CAM*), CAM2. *TCH2* and *TCH3* encode calmodulin-like (*CML*) proteins (*CML24* and *CML12*, respectively). The *TCH4* gene encodes xyloglucan endotransglycosylase/hydrolase, which is an enzyme involved in cell wall remodeling (McCormack and Braam, 2003; Toyota and Gilroy, 2013). In the genome of *Arabidopsis* 6 *CAM*, 50 *CML,* and 33 *XTH* genes have been identified (Becnel *et al*., 2006; McCormack and Braam, 2003). Touch is not the only stimulus which induces expression of these genes. The worker (Braam, 2005) described, how *TCH* expression increased in plants sprayed with a variety of hormones; *e.g*., auxin, cytokinin, and abscisic acid - and even by simply being spraying with water. Changes by cold-shock were also displayed by Polisensky and Braam (1996).

The genes *SCARECROW (SCR)* and *SHOOT GRAVITROPISM2 (SGR2)* have been identified as affecting the differentiation of both the root and the hypocotyl endodermis. Mutation in one of the *SCR* genes lead to the shoots without endodermal cell that can not respond to the gravity (Tasaka *et al*., 1999). The *SCR* gene is expressed in the cortex/endodermal initial cells and also in the endodermal cell lineage. Its tissue-specific expression is regulated at the transcriptional level. *Scr* mutant has roots without one cell layer, because assymetric division, that normally generates cortex and endodermis, is disrupted in these mutants. Also phenotype of *SCR* mutants indicates its key role in radial patterning of the root as well of the shoot (Di Laurenzio *et al*., 1996; Wysocka-Diller *et al*., 2000). *SGR2* gene is involved in the early steps of the gravitropism response and mutants of this gene also show misshapen seed and seedling (Kato *et al*., 2002). Both genes are involved in the regulatory mechanism of shoot gravitropism in *A. thaliana.* Mutation of these genes affects both inflorescence stem and hypocotyl gravitropism (Fukaki, 1996).

As mentioned previously, loss or change of gravity is a stressful factor for plants. Plants are an important part of space research, because they will be significant components of the bioregenerative life-support system on long-term space missions. In such a system plants serve as oxygen producers, food sources, and an important element in waste management. Because of these factors it is key to know how conditions in outer space affect plants growth as well as their development.

In the current study, we evaluated the influence of simulated microgravity on gene expression of *CATALASES, LEAFY COTYLEDON* and *LATE EMBRYOGENESIS ABUNDANT genes, TOUCH, SCARECROW,* and *SHOOT GRAVITROPISM2* genes in the embryos of *A. thaliana*. The transcript levels after different exposure times under microgravity were examined using quantitative real-time PCR.

## Materials and Methods

Seeds of *Arabidopsis thaliana* (cv. Columbia) were provided by the Department of Botany (UPOL, Olomouc, Czech Republic). Materials used were: MS (Murashige and Skoog, 1962), Agar (Duchefa, Haarlem, The Netherlands), and MES (Sigma-Aldrich, St. Louis, MO). The *Arabidopsis thaliana* (cv. Columbia) were grown in a greenhouse. Acquired immature siliques, about 1 cm long, were sterilized in 70% ethanol for 2 min, then in 2,5% chloramine for 8 min, washed by sterilized water three times, and finally collected and transferred to Petri dishes with MS medium (1/2 Murashige & Skoog; 0.05% MES; 1% Agar). Next, plastic Petri dishes (5.5 cm diameter) with MS medium containing about 20-30 siliques of *Arabidopsis thaliana* were fixed in the center of the inner frame of the RPM (Random Positioning Machine; Dutch Space Company). The input max. frame speed was 50 deg/sec and the input min. frame speed was 40 deg/sec. The samples were exposed to this 3D-clinostat (shown in Fig. 1) for 48, 96, and 168 hours at room temperature. The control plants (embryos) were incubated under the same conditions and for the same time as the test samples (only without the RPM).

**Fig. 1.**
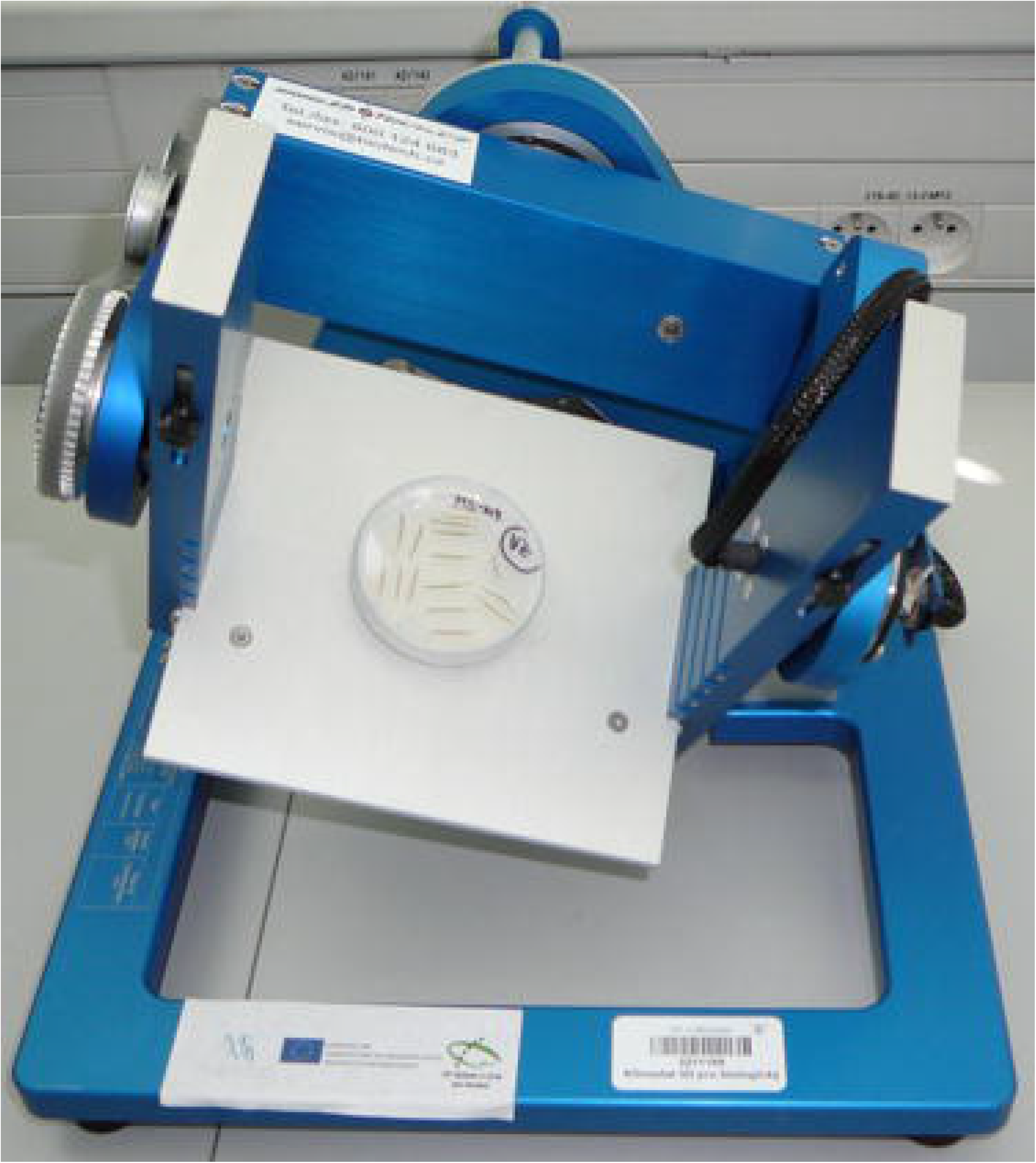
The samples of *Arabidopsis thaliana* fixed in the centre of the inner frame of the Random Positioning Machine (RPM) used in this study. RPM simulates local microgravity, thanks to the independent rotation of its two axes. This machine was developed by Japanese plant researchers and manufactured in the Netherlands (Dutch Space). The basic principle consists of an inner and an outer frame rotating independently from each other in random direction. By varying the position randomly, the RPM simulates microgravity for objects inside its container. In our case, the sample is placed in the centre of the machine.

## Quantitative reverse transcriptase polymerase chain reaction (qRT-PCR)

Total RNA was isolated from ~ 100 mg of biomass per sample of embryos after 48, 96, and 196 hours of treatment in the RPM, using a Spectrum Plant Total RNA kit (Sigma-Aldrich, St. Louis, MO, USA). The RNA was quantified by spectrophotometric analysis. Prior to amplification, all RNA samples were treated with RNase-free DNase (Promega, Madison, WI, USA) to eliminate genomic DNA contamination. The isolated RNA was transcribed to cDNA according to a common protocol by a Transcriptor High Fidelity cDNA Synthesis Kit (Roche, Basel, Switzerland) using anchored-oligo(dT)_18_ primers. The qRT-PCR was carried out on a LightCycler Nano Real - Time PCR System (Roche Diagnostic Corporation). The suitable primers (*AT2G28390, CATs, CML24*, *LEA 4-5*, *SCR*, *SGR2*) used in our study were designed with the Primer3 program, and the target sequences were obtained from the GenBank database (nucleotide database of *Arabidopsis thaliana*). The sequences for *CML12 and XTH22* were from (Fukaki et al., 1996); *LEC1, LEC2*, and *FUSCA3* were from (Kraut et al., 2011). These primers were synthesized by Generi Biotech (Hradec Králové, Czech Republic), and the appropriate primer sequences are summarized in Table 1. The qRT-PCR was performed with a LightCycler^®^ Fast Start DNA MasterPlus SYBR Green I kit (Roche Diagnostic Corporation, Prague, Czech Republic) using 10 min pre-incubation at 95°C, 40 cycles of 10 s at 95°C, 30 s at 53-64°C (temperature dependant on the respective primers), 20 s at 72°C, and 5-min incubation at 72°C. The primary data (Cq) were analyzed by LightCycler Nano Software 1.1. Gene expression was normalized per *AT2G28390* as a housekeeping gene. The data were processed by the delta-delta method (Livak and Schmittgen, 2001) and compared to the expression levels in siliques of *Arabidopsis thaliana* located in the laboratory for 48, 96, and 168 hours without RPM. All measurements were performed in triplicate. Melting curve analysis was carried out to confirm the specificity of each product. Melting curve conditions were 60°C to 97°C at 0.1°C.s^−1^. The results were analysed by 2% agarose gel electrophoresis, stained with GelRed^TM^, and photographing under UV light (254 nm) using a gel documentation system (Uvitec, Cambridge, Great Britain).

**Table 1.**
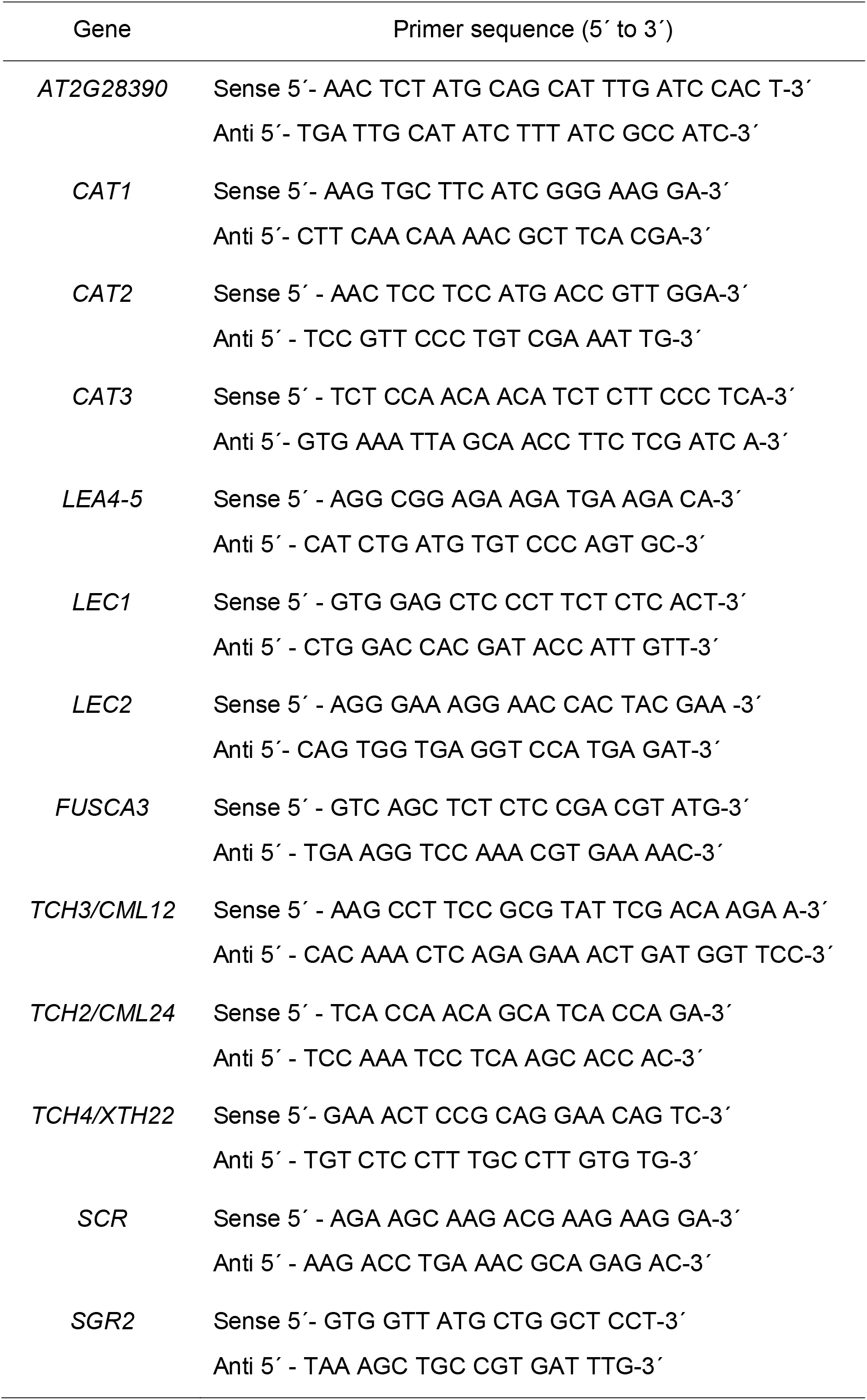
Primer sequences used in this work for qRT-PCR

## Results

In a series of experiments, the embryos of *Arabidopsis thaliana* were exposed to the simulated microgravity (RPM) for 48, 96, and 168 hours at the room temperature (Fig. 1); and the control samples were incubated under the same conditions and for the same time without RPM. In this research, we tested the relative expression of 12 genes involved in plant embryogenesis, antioxidative enzymes, mechanical stimuli, and gravitropism. After treatment under simulated microgravity, we found several genes which had increased transcript levels. First we measured levels of expression of the catalase genes (*CAT1*, *CAT2*, and *CAT3*). The highest level of relative expression was achieved for the *CAT3* gene (2.2x) after 48 hours of exposure in the RPM, followed by the *CAT2* gene (1.8x) after treatment of 168 hours. The levels of the other catalase genes were not significant. The profiles of the expression of the catalases (*CAT1*, *CAT2*, *CAT3*) of *A. thaliana* are presented in Fig. 2.

**Fig. 2.**
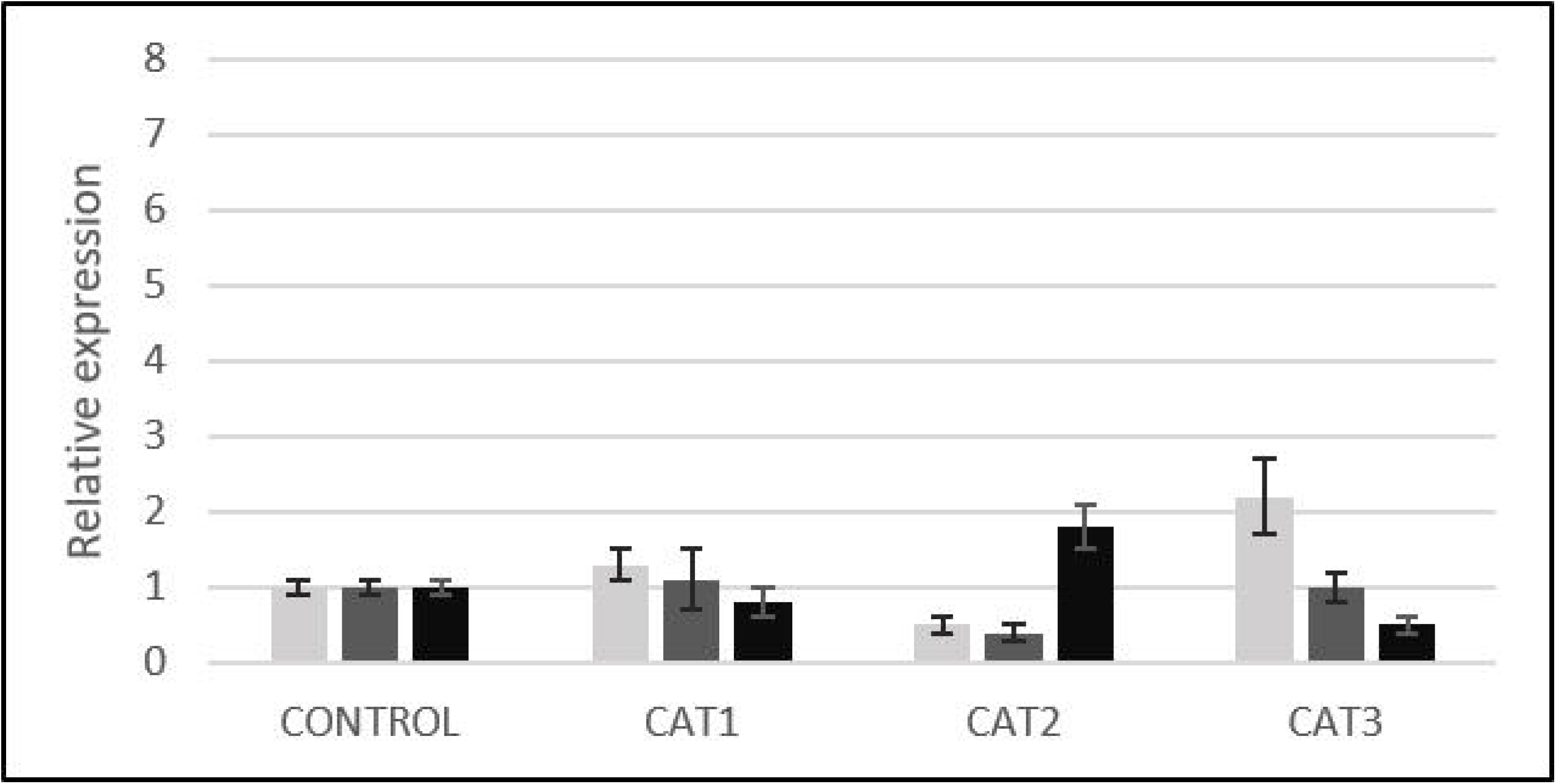
The relative expression profile of antioxidative genes (*CAT1, CAT2, CAT3*) in *Arabidopsis* embryos exposed to RPM for 48 (light grey), 96 (dark grey) and 168 (black) hours.

Second, we tested a group of genes (*LEA4-5, LEC1, LEC2, FUSCA3*) involved during embryo development (data shown in Fig. 3). For the *LEA4-5* gene, exposure to the RPM for 168 hours was the most effective, and where we determined the highest level of expression (1.7x) compared with the control samples. In the case of the *LECs* genes, significant changes in the levels of gene expression were obtained for *LEC1.* The data shows that gene expression (*LEC1*) reached the maximum level of expression after 96 hours (2.0x) of simulated microgravity. Expressions after 48 and 168 hours were at the control level; this was similar for the expression of *LEC2* and *FUSCA3*, which were also at the control level.

**Fig. 3.**
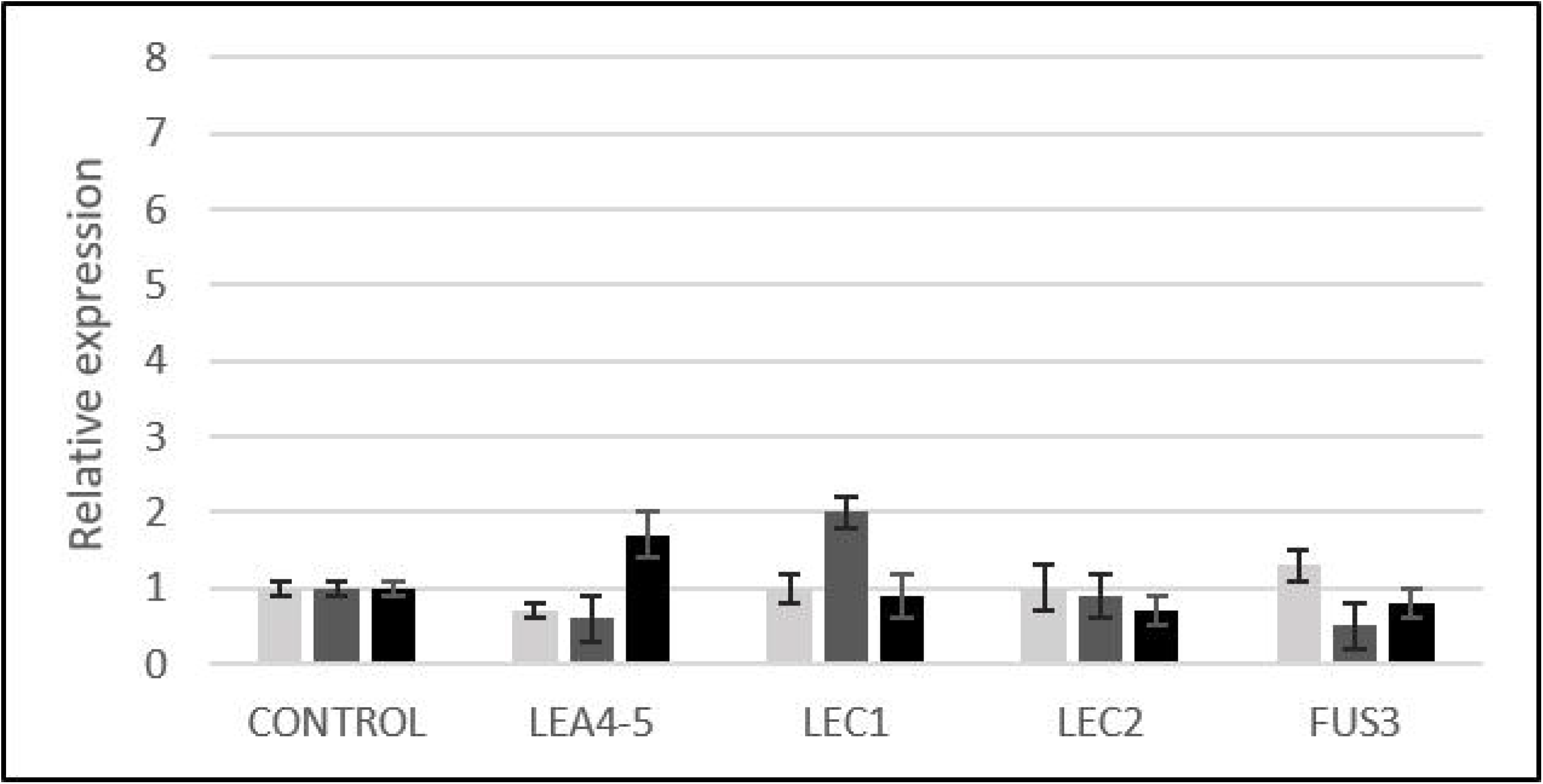
The relative expression profile of embryogenesis genes (*LEA*4-5, *LEC1*, *LEC2*, *FUSCA3*) in *Arabidopsis* embryos exposed to RPM for 48 (light grey), 96 (dark grey) and 168 (black) hours.

Third, we evaluated the levels of expression of *TOUCH* genes (*CML12*, *CML24*, *XTH22*) influenced by simulated microgravity. The best result (7.1x) was reached by the *CML24* gene after 48 hours of RPM exposure. Results for 96 and 168 hours were not significant. After 48 hours of exposure, an increase was also detected in the *XTH22* gene (1.8x); also *CML12* did not show any significant increase in expression (Fig. 4).

**Fig. 4.**
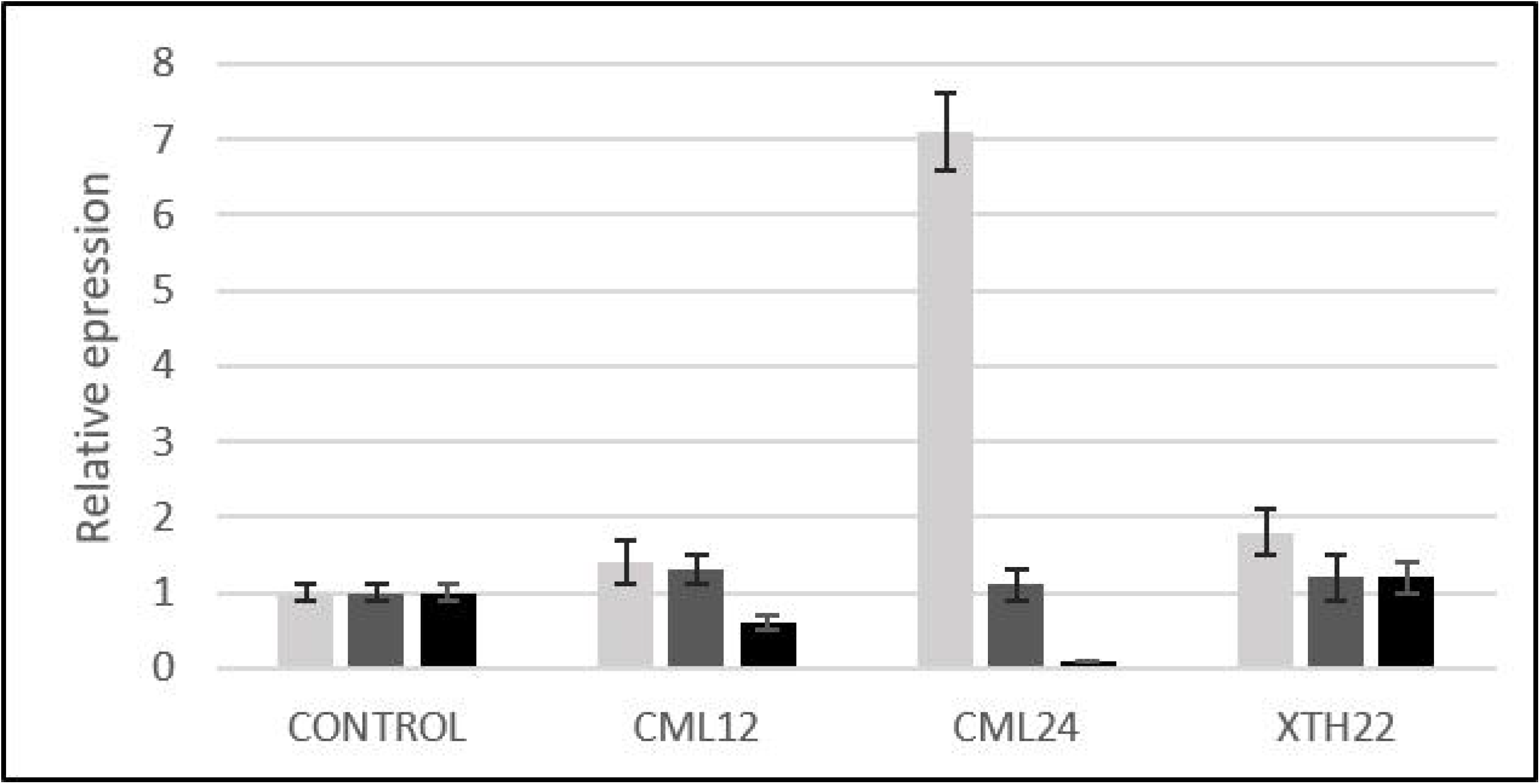
The relative expression profile of touch genes (*CML*12, *CML*24*, XTH*22) in *Arabidopsis* embryos exposed to RPM for 48 (light grey), 96 (dark grey) and 168 (black) hours.

The last interesting group of genes studied, *SCR* and *SGR2*, are genes involved in gravitropism. The most effective influence on the level of *SGR2* expression was after 168 hours of exposure under simulated microgravity. The level increased steeply after 168 hours (3.6x), in comparison with after 48 and 96 hours. Similar results were also obtained for the *SCR* gene, which showed an increased level of gene expression (1.7x), also after 168 hours of the embryos in the RPM. The other expression values for shorter times of exposure under simulated microgravity (48, 96 h) were negligible. The data obtained confirm the positive effect of simulated microgravity on genes involved in gravitropism. The relative expression profile of the gravitropism genes (*SCR, SGR2*) is summarized in Fig. 5.

**Fig. 5.**
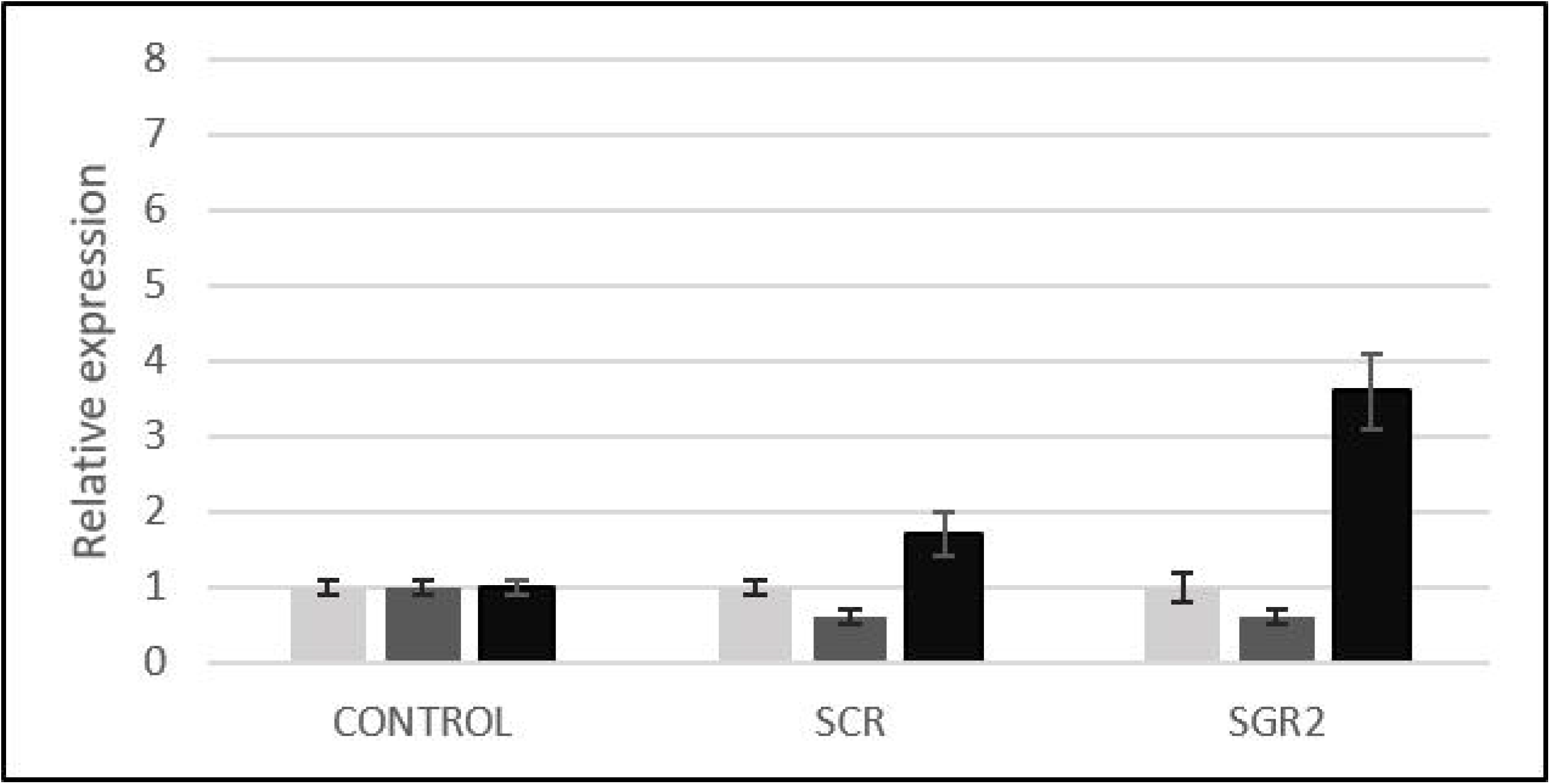
The relative expression profile of gravitropism genes (*SCR, SGR2*) in *Arabidopsis* embryos exposed to RPM for 48 (light grey), 96 (dark grey) and 168 (black) hours.

## Discussion

In this research, we studied the effects of simulated microgravity (RPM) on gene expression in *A. thaliana* over exposures times of 48, 96, and 168 hours. First, we evaluated the levels of expression of catalases in *A. thaliana* (*CAT1*, *CAT2*, and *CAT3*). It is known, in *Arabidopsis,* catalase genes encode a small family of proteins, which have an important role in controlling homeostasis of reactive oxygen species by catalyzing decomposition of H_2_O_2_ (Du *et al.*, 2008). Alam and Ghosh (2018) studied the expression of the catalase genes of *A. thaliana* and *O. sativa*. In *A. thaliana,* they observed that all three isozymes were highly expressed in the floral organs and seed. Also, they asserted that *CAT1* kept a very low level of expression in most of the studied tissues. Furthermore, all these transcripts behaved differently during various stresses (cold, drought, salt, oxidative, osmotic, and biotic treatments). The researchers Du *et al*. (2008) analyzed the expression profiles in *A. thaliana* of three catalases under different treatments such as drought, cold, oxidative stresses, salicylic acid, etc. Under all of the stress conditions, the mRNA abundance of *CAT1* was significantly increased. *CAT1* has an important role in the elimination of H_2_O_2_, generated under different environmental stresses. The drought and cold stresses activated the *CAT2* gene, and *CAT3* was mainly enhanced by oxidative treatments as well as at the senescence stage. In the study by Hausmann et *al.* (2014) callus cell cultures of *A. thaliana* (cv. Columbia) were exposed to parabolic flights in order to assess short-term molecular responses to a modified gravity field. They described that 20 seconds of microgravity, caused an immediate rise in levels of H_2_O_2_ and cytosolic Ca^2+^, and also alternation in gene expression were observed.

Second, we evaluated the levels of the expression of four genes which are involved in the process of plant embryogenesis (*LEA4-5, LEC1, LEC2*, and *FUSCA3*). *LEA4-5* is part of the fourth group of *LEA* genes. Proteins encoded by this group accumulate in dry embryos and in vegetative tissue during times of water deficit (Battaglia *et al.,* 2008). In the paper by Bies-Ethève *et al*. (2008) the expression profiles of several *LEA* genes of *A. thaliana* in wild-type background were analyzed. The *LEA4-5* gene had its strongest expression in seeds, especially in the latest stage of seed maturation (18 - 21 day after pollination). The accumulation of LEA4-5 protein in late embryogenesis and dry seed was also observed by Olvera-Carrillo *et al.* (2010). These author and co-authors (Bies-Ethève *et al*., 2008) also indicated that overproduction of LEA4-5 protein can increase the tolerance to severe drought. Another research paper (Hundertmark and Hincha, 2008) studied the expression profile of 51 *A. thaliana LEA* genes, included *LEA 4-5*, under several stress conditions (cold, drought, high light, salt, heat, mildew). In general, the greatest increases in expression were detected under cold, drought and salt conditions. In the case of *LEA4-5* gene, the expression was mainly enhanced by salt, but was also enhanced by drought conditions.

Gaj *et al*., (2005) observed a total loss of somatic embryogenesis competence in cultures of mutants with two or three mutated *LEC/FUS* genes. In the research by Ikeda-Iwai *et al*. (2002), they observed the expression of *LEC1* and *FUS3* genes in young siliques of *Arabidopsis* containing zygotic embryos, in primary and secondary somatic embryos, as well as in embryogenic callus; but not in vegetative organs such as flower buds and stems. A similar research paper (Ikeda-Iwai *et al*., 2002) concluded that *FUS3* gene expression was also observed in stress-induced somatic embryos of *Arabidopsis* produced during the culturing of shoot-apical-tips and floral buds. The *LEC1* and *LEC2* genes, having similar but not identical functions, were each sufficient to induce embryogenic competence in somatic cells.

Third, we evaluated the levels of three touch genes (*CML12/TCH3, CML24/TCH2, XTH22/TCH4*). Expression of these genes is induced by diverse mechanical stimuli, such as touch, wind, water spray, and cold (Polisensky *et* Braam, 1996; Braam *et al*., 1997) In the study by Lee *et al*. (2005) changes in the expression of *A. thaliana* genes by touch and darkness were examined using an Affymetrix Arabidopsis gene chip. They found that 589 genes were induced by touch and 461 by darkness. Over 300 genes were induced by both stimuli. Three *TCH* genes were included in this study (*CML12/TCH3, CML24/TCH2, XTH22/TCH4*), and all of them were induced by both stimuli. Ca^2+^ is probably involved in the signaling pathways of touch regulated genes (Braam, 1992; Wright *et al.*, 2002). In the study by Braam (1992) the level of extracellular Ca^2+^ was increased, which led to the induction of expression of the *TCH2, TCH3,* and *TCH4* genes. As mentioned before, 20 s of microgravity during parabolic flight increased the levels of cytosolic Ca^2+^, which could affect the expression of *TCH* genes (Hausmann *et al*., 2014).

Lastly, we measured the levels of expression of *SCR* and *SGR2* genes in *Arabidopsis.* Tasaka *et al.* (1999) described the mutation of the *SCR* and *SGR2* genes, and confirmed that after mutation of these genes the shoots did not respond to gravity; indicating that endodermal cells are essential for gravitropism of the shoot. The research paper by Wysocka-Diller *et al.* (2000) described how mutation of the *SCR* gene results in a radial pattern defect and the loss of the ground tissue layer in the root. They showed how *scr* mutants revealed that both hypocotyl and shoot inflorescence also have a radial pattern defect, loss of the normal starch sheath layer; and consequently are unable to sense gravity in the shoot. The mutation at either the *SCR* (*SGR1)* or *SGR2* locus causes a gravitropic defect in hypocotyls, this information about these genes are discussed in study (Fukaki *et al*., 1996). The comparison of a *sgr1* mutant which exhibited abnormal shoot gravitropism with wild type is presented in work (Fukaki *et al*., 1998).

Based on previous studies, plants are able to grow and reproduce in outer space conditions Nevertheless, some abnormalities were observed, including chromosomal breakage, failures in seed production, nonviable embryos, and changes in gene expression (Link *et al*., 2014).

## Conclusions

We conclude, that modeled microgravity using a RPM can alter some transcriptional activity of the genes studied in *Arabidopsis thaliana*. From our interesting results we can state that microgravity primarily alters the gene expression of *TCH2/CML24,* followed by *SGR2*, *CAT3,* and *LEC1*, respectively. Changes in gene expression were also detected in the rest of the genes, but these increases were under 2x. The effect of simulated microgravity not only depended on the specific gene, but also on the exposure time under the simulated microgravity. In the future, more studies should be performed on the effects of microgravity upon gene expression in plants, and these studies should include more than one generation of these plants.

## Acknowledgments

This research was supported by the grant IGA Prf-2018-001.

## References

Alam NB, Ghosh A. 2018. Comprehensive analysis and transcript profiling of *Arabidopsis thaliana* and *Oryza sativa* catalase gene family suggests their specific roles in development and stress responses. Plant Physiology and Biochemistry 123, 54–64.

Battaglia M, Olvera-Carrillo Y, Garciarrubio A, Campos F, Covarrubias AA. 2008. The Enigmatic LEA Proteins and Other Hydrophilins. Plant Physiology 148 (1), 6–24.

Becnel J, Natarajan M, Kipp A, Braam J. 2006. Developmental expression patterns of Arabidopsis *XTH* genes reported by transgenes and Genevestigator. Plant Molecular Biology 61 (3), 451–467.

Bies-Ethève N, Gaubier-Comella P, Debures A, Lasserre E, Jobet E, Raynal M, Cooke R, Delseny M. 2008. Inventory, evolution and expression profiling diversity of the LEA (late embryogenesis abundant) protein gene family in Arabidopsis thaliana. Plant Molecular Biology 67 (1-2), 107–124.

Borst AG, van Loon JJWA. 2009. Technology and Developments for the Random Positioning Machine, RPM. Microgravity Science and Technology 21, 287–292.

Braam J. 1992. Regulated expression of the calmodulin-related *TCH* genes in cultured *Arabidopsis* cells: induction by calcium and heat shock. Proceedings of the National Academy Sciences of the United States of America 89 (8), 3213–3216.

Braam J, Sistrunk ML, Polisensky DH, Xu W, Purugganan MM, Antosiewicz DM, Campbell P, Johnson KA. 1997. Plant responses to environmental stress: regulation and functions of the Arabidopsis *TCH* genes. Planta 203 (Suppl1), S35–S41.

Braam J. 2005. In touch: plant responses to mechanical stimuli. New Phytologist 165 (2), 373–389.

Braybrook SA, Harada JJ. 2008. LECs go crazy in embryo development. Trends in Plant Science 13 (12), 624–630.

Di Laurenzio L, Wysocka-Diller J, Malamy JE, Pysh L, Helariutta Y, Freshour G, Hahn MG, Feldmann KA, Benfey PN. 1996. The SCARECROW Gene Regulates an Asymetric Cell Division That Is Essential for Generating the Radial Organization of the Arabidopsis Root. Cell 86 (3), 423–433.

Du Y-Y, Wang P-Ch, Chen J, Song Ch-P. 2008. Comprehensive functional analysis of the catalase gene family in *Arabidopsis thaliana*. Journal of Integrative Plant Biology 50 (10), 1318–1326.

Frugoli JA, Zhong HH, Nuccio ML, McCourt P, McPeek MA, Thomas TL, McClung CR. 1996. Catalase is encoded by a multigene family in Arabidopsis thaliana (L.) Heynh. Plant Physiology 112 (1), 327–336.

Fukaki H, Fujisawa H, Tasaka M. 1996. *SGR1, SGR2,* and *SGR3:* Novel Genetic Loci Involved in Shoot Gravitropism in *Arabidopsis thaliana*. Plant Physiology 110 (3), 945–955.

Fukaki H, Wysocka-Diller J, Kato T, Fujisawa H, Benfey PN, Tasaka M. 1998. Genetic evidence that the endodermis is essential for shoot gravitropism in *Arabidopsis thaliana*. The Plant Journal 14 (4), 425–430.

Gaj MD, Zhang S, Harada JJ. Lemaux PG. 2005. Leafy cotyledon genes are essential for induction of somatic embryogenesis of *Arabidopsis*. Planta 222, 977–988.

Harada JJ. 2001. Role of *Arabidopsis LEAFY COTYLEDON* gene in seed development. Journal of Plant Physiology 158, 405–409.

Hausmann N, Fengler S, Hennig A, Franz-Wachtel M, Hampp R, Neef M. 2014. Cytosolic calcium, hydrogen peroxide and related gene expression and protein modulation in *Arabidopsis thaliana* cell cultures respond immediately to altered gravitation: parabolic flight data. Plant Biology 16, 120–128.

Hundertmark M, Hincha DK. 2008. LEA (Late Embryogenesis Abundant) proteins and their encoding genes in *Arabidopsis thaliana*. BMC Genomics 9, 118.

Ikeda-Iwai M, Satoh S, Kamada H. 2002. Establishment of a reproducible tissue culture system for the induction of Arabidopsis somatic embryos. Journal of Experimental Botany 53 (374), 1575-1580.

Islam T, Manna M, Kaul T, Pandey S, Reddy CS, Reddy M. 2015. Genome-wide dissection of *arabidopsis* and rice for the identification and expression analysis of glutathione peroxidases reveals their stress-specific and overlapping response patterns. Plant Molecular Biology Reporter 33 (5), 1413–1427.

Kato T, Morita MT, Fukaki H, Yamauchi Y, Uehara M, Niihama M, Tasaka M. 2002. SGR2, a Phospholipase-Like protein, and ZIG/SGR4, a SNARE, are involved in the shoot gravitropism of Arabidopsis. Plant Cell 14 (1), 33–46.

Kraut M, Wójcikowska B, Ledvon A, Gaj MD. 2011. Immature Zygotic Embryo Cultures of *Arabidopsis*. A Model System for Molecular Studies on Morphogenic Pathways Induced In Vitro. Acta Biologica Cracoviensia Series Botanica 52/2, 59–67.

Lee D, Polisensky DH, Braam J. 2005. Genome-wide identigication of touch- and darkness-regulated Arabidopsis genes: a focus on calmoduline-like and XTH genes. The New Phytologist 165 (2), 429–444.

Link BM, Busse JS, Stankovic B. 2014. Seed-to-seed-to-Seed Growth and Development of *Arabidopsis* in Microgravity. Astrobiology 14 (10), 866–875.

Livak KJ, Schmittgen TD. 2001. Analysis of relative gene expression data using real-time quantitative PCR and the 2(-Delta Delta C(T)) Method. Methods 25 (4), 402–408.

Martzivanou M, Babbick M, Cogoli-Greuter M, Hampp R. 2006. Microgravity-related changes in gene expression after short-term exposure of *Arabidopsis thaliana* cell cultures. Protoplasma 229 (2-4), 155–162.

McClung CR. 1997. Regulation of Catalases in Arabidopsis. Free Radical Biology and Medicine 23 (3), 489–496.

McCormack E, Braam J. 2003. Calmodulins and related potential calcium sensors of Arabidopsis. New Phytologist 159 (3), 585–598.

Mhamdi A, Queval G, Chaouch S, Vanderauwera S, Van Breusegem F, Noctor G. 2010. Catalase function in plants: a focus on Arabidopsis mutants as stress-mimic models. Journal of Experimental Botany 61 (15), 4197–4220.

Olvera-Carrillo Y, Campos F, Reyes JL, Garciarrubio A, Covarrubias AA. 2010. Functional Analysis of the Group 4 Late Embryogenesis Abundant Protein Reveals Their Relevance in the Adaptive Response during Water Deficit in Arabidopsis. Plant Physiology 154 (1), 373–390.

Polisensky DH, Braam J. 1996. Cold-shock regulation of the Arabidopsis *TCH* genes and the effects of modulation intracellular calcium levels. Plant Physiology 111, 1271–1279.

Stone SL, Kwong LW, Yee KM, Pelletier J, Lepiniec L, Fischer RL, Goldberg RB, Harada JJ. 2001. *LEAFY COTYLEDON2* encodes a B3 domain transcription factor that includes embryo development. Proceedings of the National Academy Sciences of the United States of America 98 (20), 11806-11811.

Tasaka M, Kato T, Fukaki H. 1999. The endodermis and shoot gravitropism. Trends in Plant Science 4 (3), 103–107.

Toyota M, Gilroy S. 2013. Gravitropism and mechanical signalling in plants. American Journal of Botany 100 (1), 111–125.

West MAI, Yee KM, Danao J, Zimmerman JL, Fischer RL, Goldberg RB, Harada JJ. 1994. *LEAFY COTYLEDONl* 1s an Essential Regulator of Late Embryogenesis and Cotyledon ldentity in-Arabidopsis. The Plant Cell 6, 1731–1745.

Wright AJ, Knight H, Knight MR. 2002. Mechanically stimulated *TCH3* gene expression in Arabidopsis involves protein phosphorylation and EIN6 downstream of calcium. Plant Physiology 128 (4), 1402–1409.

Wysocka-Diller JW, Helariutta Y, Fukaki H, Malamy JE, Benfey PN. 2000. Molecular analysis of SCARECROW function reveals a radial patterning mechanism common to root and shoot. Development 127 (3), 595–603.

Zimmermann P, Heinlein C, Orendi G, Zentgraf U. 2006. Senescence-specific regulation of catalases in *Arabidopsis thaliana* (L.) Heynh. Plant Cell & Environment 29 (6), 1049–1060.

